## Supplementary Information for "Optogenetic stimulation of anterior insular cortex neurons reveals causal mechanisms underlying suppression of the default mode network by the salience network"

**Dynamic decoupling of salience and default mode networks by optogenetic manipulation of anterior insular cortex**

**Supplementary Information**

**I. Supplementary Materials and Methods ---------------------------------------** Page 2

**II. Supplementary Results ----------------------------------------------------------** Page 5

**III. Supplementary Discussion -----------------------------------------------------** Page 12

**IV. Supplementary Figures ---------------------------------------------------------** Page 14

**I. Supplementary Materials and Methods**

**1. Bayesian Switching Dynamic Systems model**

To determine latent brain state dynamics underlying optogenetic manipulation of the anterior insular cortex (AI), we used a Bayesian Switching Dynamic Systems (BSDS) model (1). Here we briefly describe the mathematical framework of the BSDS model(1). Let $\boldsymbol{y}_{t}^{s}$ denote a D-dimensional vector of ROI timeseries obtained from subject s in time t, where D is the number of ROIs. Following the general formulation of the switching state-space models, we defined $\boldsymbol{z}_{t}^{s}$ as the latent state variables and $\boldsymbol{x}_{kt}^{s}$ as the latent space variables associated to $\boldsymbol{y}_{t}^{s}$ at the $k$-th latent state, that is $z_{kt}^{s}=1$. The $\boldsymbol{z}_{t}^{s}$ is a 1-of-$K$ discrete vector with elements $z_{kt}^{s}$, $\forall k=1,\ldots,K$. Two successive time instances are dependent through a 1st-order Markov chain of Hidden Markov Model (HMM). Using Markovian properties and given state transition probabilities $\boldsymbol{A}$, where $A_{jk}\equiv p\left( z_{kt}^{s}=1 \right|\boldsymbol{z}_{j,t-1}^{s}=1)$ and a marginal distribution $p\left( \boldsymbol{z}_{1}^{s} \right|\boldsymbol{\pi})= \prod_{k=1}^{K} \pi_{k}^{z_{k1}^{s}}$ represented by a vector of initial probabilities $\boldsymbol{\pi}$ where $\pi_{k}\equiv p(z_{k1}^{s}=1)$, the probability distribution for the latent state variables is expressed by $p\left( \boldsymbol{z}_{t}^{s} \right|\boldsymbol{z}_{t-1}^{s}, \boldsymbol{A})=\prod_{k=1}^{K} \prod_{j=1}^{K} A_{jk}^{z_{j,t-1}^{s} z_{kt}^{s}}$ for all $t>1$. We assume that at a given latent state $k$ in time $t$, shown by $z_{kt}^{s}=1$, the observed vector $\boldsymbol{y}_{t}^{s}$ is generated via probabilistic interpretation of a factor analysis model (2, 3) as:

$\boldsymbol{y}_{t}^{s}=\boldsymbol{U}_{k}\boldsymbol{x}_{kt}^{s} +\boldsymbol{\mu}_{k}+\boldsymbol{e}_{kt}$, $\forall t |z_{kt}^{s}=1$,

where $\boldsymbol{U}_{k}$ is a $D\times P$ dimensional linear transformation matrix that transforms data to a subspace of lower dimensionality, $P<D$, described using a $P$-dimensional vector of latent space variables $\boldsymbol{x}_{kt}^{s}$ mediated by an overall bias $\boldsymbol{\mu}_{k}$ and a measurement noise $\boldsymbol{e}_{kt}$. With the normality assumption, that is $\boldsymbol{x}_{kt}^{s}\mathcal{\sim N}\left( \mathbf{0}, I \right)$ and $\boldsymbol{e}_{kt}\mathcal{\sim N}\left( \mathbf{0},\boldsymbol{\Psi}_{k} \right)$, the marginal distribution of $\boldsymbol{y}_{t}^{s}$ follows a Gaussian distribution as $p\left( \boldsymbol{y}_{t}^{s} \right|\boldsymbol{\mu}_{k}, \boldsymbol{U}_{k}, \boldsymbol{\Psi}_{k}\mathcal{)=N}\left( \boldsymbol{\mu}_{k}, \boldsymbol{U}_{k}{\boldsymbol{U}_{k}}^{T}+\boldsymbol{\Psi}_{k} \right)$ where $T$ denotes the transpose operator. We then define a dynamical process on the latent space variables using an autoregressive (AR) model (4) of order $R$ as:

$\boldsymbol{x}_{kt}^{s}={\bar{\boldsymbol{X}}}_{kt}^{s}\vec{V_{k}} +\boldsymbol{\varepsilon}_{kt}$, $\forall t |z_{kt}^{s}=1$,

where $\vec{V_{k}}$ is a vector of AR coefficients. ${\bar{\boldsymbol{X}}}_{kt}^{s}=diag({\bar{\boldsymbol{x}}}_{kt}^{s}$) is a block diagonal isotropic matrix with elements of ${\bar{\boldsymbol{x}}}_{kt}^{s}=\left( {{\bar{\boldsymbol{x}}}_{k,t-1}^{s}}^{T}, {{\bar{\boldsymbol{x}}}_{k,t-2}^{s}}^{T}, \ldots, {{\bar{\boldsymbol{x}}}_{k,t-R}^{s}}^{T} \right)$ represented using latent space variables from the previous $R$ time frames. $\boldsymbol{\varepsilon}_{kt}\mathcal{\sim N}\left( \mathbf{m}_{k},\boldsymbol{\Sigma}_{k} \right)$ is the remaining error term in the latent space. All analyses conducted in this study use a 1st-order AR model ($R=1$). Detailed theoretical derivations are provided in the previous study (1).

We used a variational Bayesian (VB) framework to infer model parameters, including the number of brain states. The number of states is treated as a random variable, whose optimal value is learned from data using automatic relevance determination procedures implemented in a variational Bayesian framework (1). BSDS models were initialized with 10 states for both Chronos and EYFP rats. BSDS identified 5 latent brain states in the Chronos rats and 3 latent brain states in the EYFP rats.

**2. Temporal properties of latent brain states**

BSDS estimated the posterior probability of each latent brain states at each time point and chose the latent brain state with the highest probability as the dominant state at that time point. Using the temporal evolution of the latent brain states, we measured temporal properties of each latent brain state, including occupancy rate and state switching probability. Occupancy rate quantifies the proportion of time that a state is chosen as the dominant state. State switching probability quantifies the chance that brain state at time point *t* either remains at its own state or switch to another brain state at the time point *t+1*. These temporal properties were examined to characterize their relationship with optogenetic stimulation conditions.

**3. Stimulation block prediction using time-varying latent brain state dynamics**

Stimulation block classification analysis was performed to investigate whether time-varying latent brain state dynamics contain information associated with stimulation OFF block and stimulation ON block. We built a multiclass classifier based on a linear support vector machine using the MATLAB package LIBSVM (5) to discriminate stimulation OFF block and stimulation ON block at each time point. The posterior probabilities of the latent brain states at each time point during stimulation OFF blocks and stimulation ON blocks were used as features to train the classifier. Classifier performance was evaluated by conducting leave-one-out cross-validation (LOOCV) analysis. Specifically, posterior probability time-series of the five latent brain states from one rat were used as a test set and the posterior probability time-series of the brain states obtained from rest of the rats were used to train the classifier. Then, the trained classifier was applied to the test set to predict moment-by-moment correspondence between posterior probabilities of the latent brain states and stimulation block (OFF/ON). This procedure was performed *S* times (*S*: number of rats), and the cross-validation accuracy across the test sets was used to evaluate the performance of the classifier. We further evaluated statistical significance of LOOCV accuracy using permutation testing (500 times). In each permutation, ON and OFF block labels were randomly permuted. We followed the same LOOCV analysis procedure described above to evaluate the classification accuracy under permutation. We repeated this procedure 500 times, and used the resulting distribution to evaluate the statistical significance of the LOOCV accuracy.

**4. Regional activation and functional connectivity of latent brain states**

Each latent brain state is represented by a multivariate Gaussian distribution, which is described by the mean (activation levels of ROIs) and covariance (functional connectivity between ROIs) matrices. To investigate changes in activation level of each ROI between different latent brain states, we conducted paired *t*-tests on the rat-state-wise mean values for each ROI. To determine which dynamic functional connections are important for distinguishing different latent brain states, we first computed the partial correlations from the estimated covariance matrices for each state and then conducted paired t-tests on the rat-state-wise z-transformed correlation matrices. Multiple comparisons were corrected using false discovery rates (*p* < 0.05).

**5. General linear model (GLM) analysis of brain activation and connectivity associated with AI stimulation**

We used a conventional general linear model (3dDeconvolve) as implemented in AFNI (6) to determine brain activation associated with AI stimulation. In each rat, a regressor of interest corresponding to the ON stimulation condition was convolved with a negative monocrystalline iron oxide nanoparticle (MION) hemodynamic response function (HRF)(7). Contrast β-maps corresponding to the stimulation ON condition were determined in each rat, and entered into a group level analysis. A one sample *t*-test (3dttest++) was used to determine group activation maps, and a voxel-wise threshold of *p* < 0.005 with family-wise cluster-correction threshold of *p* < 0.01 (cluster size = 38 voxels) was used to determine significant activation clusters.

We used a generalized psychophysiological interaction (gPPI) (8) model to determine changes in connectivity associated with AI stimulation. The gPPI model consisted of physiological terms, psychological terms, and PPI terms. The physiological terms were time-series data from the AI ROI; the psychological terms were the convolutions of the MION HRF with the main task effects of interest (ON condition timestamps); and the PPI terms were deconvolved raw time-series data from the AI ROI multiplied by the main effect of interest followed by a convolution with the MION HRF.

**II. Supplementary Results**

**1. GLM-based activation and connectivity associated with AI stimulation**

**1.1. Activation:** GLM analysis uncovered strong responses in the AI stimulation site in Chronos rats (**Figure S1**) and deactivation of the striatum and medial temporal lobe, but no response clusters in other brain regions (*p* < 0.005 voxel-wise threshold, cluster threshold *p* < 0.01). Inspection of the time-series from our *a priori* ROIs suggest a complex profile of temporal changes elicited by optogenetic stimulation (**Figure S2**). No significant clusters were detected in EYFP rats.

**1.2. gPPI connectivity:** gPPI connectivity analysis revealed no significant clusters (*p* < 0.005 voxel-wise threshold, cluster threshold *p* < 0.01) associated with AI stimulation in either Chronos or EYFP rats.

These results highlight the ability of latent space dynamical switching models like BSDS to capture dynamic circuit properties that are missed by conventional general linear models.

**2. Occupancy rates of latent brain states within AI stimulation ON and OFF sub-blocks**

To further characterize stimulation-dependent changes in occupancy rates of latent brain states, we divided the stimulation OFF and ON blocks into 10 s sub-blocks (**Figure S3A, B**) in Chronos rats and EYFP controls, and examined occupancy rates of the brain states in each sub-block.

In Chronos rats, we found that the occupancy rate of the ON state (State 2) was significantly higher than other brain states in both sub-blocks of the stimulation ON block (**Figure S3C, left**, *p* < 0.05, two-tailed *t*-test, FDR-corrected), suggesting that the ON state dominates the stimulation ON block. Furthermore, the occupancy rate of the ON state (State 2) was significantly higher than other brain states in the 1^st^ sub-block of the stimulation OFF block, which occurs immediately after the stimulation ON block (**Figure S3C, right**), suggesting that fMRI responses evoked by the stimulation can persist for at least 10 s. Subsequently, from the 1^st^ to 3^rd^ sub-block of the stimulation OFF block, the occurrence rate of the ON state progressively decreased, while the occupancy rate of State 3 and the OFF state gradually increased with time, eventually leading to sub-blocks dominated by the OFF state (**Figure S3C, right**). These findings suggest that State 3 may serve as an intermediate state that connects the ON and OFF states during stimulation ON-OFF boundaries and share some functional properties of the OFF state. The OFF state (State 1) dominated the remainder of the OFF block, from sub-blocks 4 through 8, suggesting that State 1 represents the latent brain state dynamic for stimulation OFF periods (**Figure S3C, left**, right, *p* < 0.05, two-tailed *t*-test, FDR-corrected). Next, analysis of EYFP controls revealed that a single state (i.e., State 1) dominated all the sub-blocks during the stimulation ON and OFF periods (**Figure S3D**, *p* < 0.05, two-tailed *t*-test, FDR-corrected), confirming that latent brain state dynamics are related to the optogenetic stimulation of AI and not induced by potential nonspecific effects of the stimulation paradigm.

Together, these analyses demonstrate that, in Chronos rats, State 1 (the OFF state) represents baseline activity, and State 2 (the ON state) represents the stimulation-evoked activity, and lastly, State 3 shares some functional similarity to the OFF state and participates as an intermediate state that connects the ON and OFF states during stimulation ON-OFF boundaries. Importantly, these results highlight the ability of latent space dynamical switching models like BSDS to capture dynamic circuit properties that would be missed by conventional GLM approaches.

**3. Dynamic causal relationships between temporal dynamics of latent brain states**

We examined how latent brain states influence each other. To accomplish this, we examined dynamic causal relations between time-varying posterior probabilities of the latent brain states as estimated by BSDS (**Figure 3A**). As noted, each latent brain state is characterized by a unique pattern of dynamic functional connectivity that transiently links distributed brain regions; investigation of causal interactions between latent brain states could provide additional insights into dynamic brain circuit mechanisms underlying interactions between the ON and OFF states.

Analysis of causal relationships between latent brain states using multivariate dynamic state-space systems identification (9-12), revealed that, during the stimulation OFF block, the OFF state had a negative causal influence on the ON state (**Figure S4A**, *p* < 0.05, two-tailed *t*-test, FDR-corrected). This pattern was reversed during the stimulation ON block, during which the ON state had a negative causal influence on the OFF state (**Figure S4B**, *p* < 0.05, two-tailed *t*-test, FDR-corrected). Furthermore, changes in posterior probability of the ON state were significantly correlated with the temporal profile of the AI response (**Figure S4C**, *r* = 0.62, *p* = 7.1x10^-9^, two-tailed *t*-test), suggesting that stimulation induces activation of the ON state and suppression of the OFF state (**Figure S4D**). These results further demonstrate the mutually inhibitory influence of brain states associated with AI engagement and retrosplenial cortex (RSC) suppression.

**4. Inter-regional functional connectivity using time-series data from BSDS-derived ON and OFF states**

We conducted additional analyses to validate inter-regional functional connectivity estimated by the BSDS model. We used data from time-points dominated by ON and OFF states, and computed partial correlations across ROIs (**Figure S5**). We specifically examined stimulation-related functional connectivity changes of the AI, PrL, and posterior RSC (-6.86 mm AP) with all other ROIs based on results shown in **Figure 4D-F**. For time-points dominated by the ON state, we found decreased connectivity between AI and anterior RSC (-2.90 mm AP), PrL and posterior RSC (-7.82 mm AP), and between RSC regions (-6.86 mm AP to -2.90 and -3.86 mm AP); we also found increased connectivity between PrL and Cg (**Figure S6A-C**). Thus, direct estimation of functional connectivity using data from time-points corresponding to the ON and OFF states yielded convergent results and validated findings from the BSDS-estimated covariance matrices.

**5. Dynamic state transition properties at stimulation boundary and OFF periods**

We examined dynamic state transition properties using the BSDS-derived state-switching matrices of each rat. Because the duration of OFF-stimulation blocks is relatively long compared to the duration of the ON block, it is possible that state transition properties during OFF-stimulation blocks could smear the state transition properties at stimulation boundaries. To address this, we computed state-switching probability matrices separately for 3 time periods: (1) 60 s beginning 10 s into OFF-stimulation blocks, (2) 20 s OFF🡪ON stimulation boundaries beginning 10 s before ON-stimulation blocks, and (3) 20 s ON🡪OFF stimulation boundaries beginning 10 s before OFF-stimulation blocks (**Figure S7A**).

Analysis of the state switching matrix during OFF-stimulation periods revealed that State 1 (the OFF state) and State 3 have higher probabilities of switching between each other than to other states (**Figure S7C, F**), suggesting that State 3 may have a role as an additional OFF state. Analysis of state switching matrices for OFF🡪ON and ON🡪OFF stimulation boundaries revealed that the transition path between OFF and ON states is most likely to include State 3 (P_OFF🡪State3_ = 0.28, P_State3🡪ON_ = 0.20, P_ON🡪State3_=0.11, P_State3🡪OFF_=0.18, **Figure S7D, E, G, H**), suggesting a role for State 3 as a Transition state during stimulation boundaries. State 3 also has the second-highest occupancy rate during OFF-stimulation blocks (**Figure S7B**), consistent with the observation that this state functions as both a secondary OFF state and an intermediate state between ON and OFF states.

Lastly, examination of temporal properties of State 4 and 5 showed that they not only have low occurrence during stimulation boundaries, but also do not have distinct patterns in their occurrence, suggesting that these two states do not function as Transition or OFF states.

**6. Replication of dynamic functional connectivity changes during the Transition compared to the ON and OFF states**

We extended the above analyses to validate inter-regional functional connectivity during the Transition compared to the ON and OFF states estimated by the BSDS model. We used data from time-points dominated by Transition, ON and OFF states, and computed partial correlations across ROIs (**Figure S5**). We then compared the results with findings from state covariance matrices driven directly by BSDS (**Figure 6C, F**). The two approaches for estimating connectivity changes between Transition and OFF state showed convergent results. Specifically, for time-points dominated by the Transition, compared to the OFF, state we found stronger functional connectivity between the AI and mid-RSC (-5.90 mm AP), but reduced intra-RSC connectivity between anterior (-2.90 and –3.86 mm AP) and mid-posterior subdivisions of RSC (-5.90 mm, -6.86 mm AP) (**Figure S8A**, all *ps* < 0.05, two-tailed t-test, FDR corrected).

Similarly, the two approaches also showed convergent results for connectivity changes between the ON and Transition states. The ON state showed increased connectivity between the PrL and Cg, but decreased connectivity between anterior-middle RSC (-3.86 mm, -4.86 mm AP) and posterior RSC subdivisions (-6.86 mm, -7.82 mm AP) (**Figure S8B**, all *ps* < 0.05, two-tailed *t*-test, FDR-corrected). Thus, direct estimation of functional connectivity using data from time-points corresponding to the Transition, ON and OFF states yielded convergent results and validated findings from the BSDS-estimated covariance matrices.

**7. Replication of findings using extended SN and DMN ROIs**

We examined the robustness of our findings with respect to ROI selection with additional subcortical nodes. We conducted additional analysis by incorporating hippocampus and amygdala nodes, which are known to be part of SN (13) and DMN (14), respectively (**Figure S9A**). As described below, all major findings were replicated with this new set of ROIs.

**7.1. Matching BSDS states estimated from different ROI sets:** BSDS identified 3 latent brain states. To determine whether brain states identified in the original model matched brain states identified in the extended ROI set, we conducted cross-model brain state correlation analysis. By computing Pearson’s correlation of posterior probabilities of latent brain states estimated from the two distinct models, we found high one-to-one mapping between S1_original_ and S1_extended_, between S2_original_ and S2_extended_, and between S3_original_ and S3_extended_, respectively (**Figure S9B**). These supplementary results demonstrate robustness of our main finding that distinct latent brain state dynamics dominate stimulation OFF and ON blocks, as well as their boundaries.

**7.2. Spatiotemporal properties of extended DMN and SN ROI-derived latent brain states corresponding to stimulation ON and OFF blocks:** Each latent brain state showed distinct moment-by-moment changes in posterior probability across stimulation ON and OFF blocks (**Figure S9C**). Examination of AI stimulation effects on the temporal properties of each state revealed that State 1 has a significantly higher occupancy rate than other states during stimulation OFF blocks (**Figure S9D**, all *ps* < 0.05, two-tailed *t*-test, FDR-corrected). Furthermore, the occupancy rate of State 1 was significantly higher during stimulation OFF blocks compared to stimulation ON blocks (**Figure S9D**, all *ps* < 0.05, two-tailed *t*-test, FDR-corrected), suggesting that State 1 is the primary state associated with the stimulation OFF blocks (OFF state). In contrast, State 2 had a significantly higher occupancy rate than other states during stimulation ON blocks (**Figure S9D**, all *ps* < 0.05, two-tailed *t*-test, FDR-corrected). Furthermore, the occupancy rate of State 2 was significantly higher during stimulation ON compared to stimulation OFF blocks (**Figure S9D**, all *ps* < 0.05, two-tailed *t*-test, FDR-corrected), implying that State 2 is a dominant state associated with the stimulation ON block (ON state).

We next examined dynamic functional connectivity associated with the ON and OFF brain states. Univariate link-specific analysis was conducted to determine unique functional connectivity patterns that differentiate the ON and OFF states (**Figure S9E**). This analysis revealed that the ON state has unique connectivity patterns compared to the OFF state (all *ps* < 0.01, two-tailed *t*-test, FDR-corrected). Notably, the ON state showed decreased connectivity between AI and an anterior RSC subdivision (-2.90 mm AP), suggesting decoupling between SN and DMN. In addition, the ON state showed increased connectivity between the prelimbic cortex (PrL) and cingulate cortex (Cg), and the decreased connectivity between the PrL and posterior RSC subdivisions (-6.86 mm, -7.82 mm AP) and between anterior and posterior RSC subdivisions. Furthermore, the ON state also showed increased connectivity between the amygdala and PrL, AI, and Cg. These results suggest functional involvement of the PrL, Cg, and amygdala in SN, and functional heterogeneity within the RSC underlying SN-DMN dynamics.

Taken together, these results are consistent with our main findings, highlighting the robustness of our findings.

**8. Replication of findings using SN and DMN nodes and additional striatum-MTL ROIs**

We further examined the robustness of our findings with respect to ROI selection by including nodes for the most significant response clusters detected by GLM (**Figure S1**), the striatum and the medial temporal lobe (MTL) (**Figure S10A**). Although the involvement of striatum and MTL is not as well established within large-scale resting-state networks in the rodent brain as our *a priori* ROIs, they are generally considered part of SN (15, 16) and DMN (17-19), respectively. As described in detail below, all major findings were replicated with this new set of ROIs.

**8.1. Matching BSDS states estimated from different ROI sets:** BSDS identified four latent brain states. To determine whether brain states identified in the original model matched brain states identified in the extended ROI set, we conducted cross-model brain state correlation analysis. By computing Pearson’s correlation of posterior probabilities of latent brain states estimated from the two distinct models, we found high one-to-one mapping between S1_original_ and S1_extended_, between S2_original_ and S2_extended_, and between S3_original_ and S3_extended_, respectively (**Figure S10B**). These supplementary results demonstrate robustness of our main finding that distinct latent brain state dynamics are associated with optogenetic stimulation of the AI.

**8.2. Spatiotemporal properties of extended ROI-derived latent brain states corresponding to stimulation ON and OFF blocks:** Each latent brain state showed distinct moment-by-moment changes in posterior probability across stimulation ON and OFF blocks (**Figure S10C**). Examination of AI stimulation effects on the temporal properties of each state revealed that State 1 has a significantly higher occupancy rate than other states during stimulation OFF blocks (**Figure S10D**, all *ps* < 0.05, two-tailed *t*-test, FDR-corrected). Furthermore, the occupancy rate of State 1 was significantly higher during stimulation OFF blocks compared to stimulation ON blocks (**Figure S10D**, all *ps* < 0.05, two-tailed *t*-test, FDR-corrected), suggesting that State 1 is the primary state associated with the stimulation OFF blocks (i.e., OFF state). In contrast, State 2 had a significantly higher occupancy rate than other states during stimulation ON blocks (**Figure S10D**, all *ps* < 0.05, two-tailed *t*-test, FDR-corrected). Furthermore, the occupancy rate of State 2 was significantly higher during stimulation ON compared to stimulation OFF blocks (**Figure S10D**, all *ps* < 0.05, two-tailed *t*-test, FDR-corrected), implying that State 2 is a dominant state associated with the stimulation ON blocks (i.e., ON state).

We next examined dynamic functional connectivity associated with the ON and OFF brain states. Univariate link-specific analysis was conducted to determine unique functional connectivity patterns that differentiate the ON and OFF states (**Figure S10E**). This analysis revealed that the ON state has unique connectivity patterns compared to the OFF state (all *ps* < 0.05, two-tailed *t*-test, FDR-corrected). Notably, the ON state showed decreased connectivity of the AI with anterior RSC subdivision (-2.90 mm AP), Striatum and MTL. In addition, the ON state showed increased connectivity between the PrL and Cg, and decreased connectivity between the PrL and posterior RSC subdivisions (-6.86 mm, -7.82 mm AP) and between anterior and posterior RSC subdivisions. These results suggest that the striatum and MTL are involved in SN-DMN dynamics, and are overall consistent with our main results, highlighting the robustness of our findings.

**9. Control analysis using ROIs located outside the SN and DMN**

We conducted additional control analyses using ROIs from the auditory, visual, and motor cortex outside of the canonical SN and DMN (**Figure S11A**) to investigate the specificity of our findings with respect to SN and DMN nodes. BSDS identified 4 latent brain states from the new ROI set. In contrast to our main analysis, latent brain states estimated from the new ROI set did not show stimulation-dependent changes in posterior probability (**Figure S11B**). Furthermore, examination of the occupancy rate of each latent brain state during stimulation OFF and ON blocks demonstrated that the major findings reported in our main analyses (i.e., stimulation-dependent changes in occupancy rate of latent brain states) are not observed with the new ROI set (**Figure S11C**). Taken together, these results demonstrate the specificity of our findings with respect to canonical SN-DMN ROIs.

**III. Supplementary Discussion**

**1. Rationale for anatomically-defined SN and DMN ROIs**

Because the focus of our study was to investigate AI stimulation-induced changes in SN-DMN interactions in rodent brain, we selected AI, PrL, Cg, and RSC as the most logical ROIs to conduct the analyses based on a wide range of published studies (13, 14, 16, 20-27). Because conventional GLM analyses did not uncover strong responses in these ROIs, except for the AI stimulation site (**Figures S1 and S2**), we used anatomically-defined canonical salience (SN) and default mode network (DMN) nodes encompassing the AI, PrL, Cg, and RSC (13, 14, 16, 20-27).

Our use of anatomically-defined ROIs was also motivated by inconsistencies in identification of SN and DMN nodes in resting-state fMRI studies. While analysis of functional connectivity using resting-state fMRI has identified a rodent DMN anchored in the RSC (13, 14), there is less agreement about inclusion of medial prefrontal cortex DMN nodes, such as the Cg and PrL (14, 21, 23, 24, 26, 27). Indeed, in a recent study characterizing the rodent SN, Cg and PrL were considered as key nodes of the SN in addition to their putative roles in the DMN (13). Recent imaging studies have also paradoxically assigned individual subdivisions of the RSC and medial prefrontal cortex to both the SN and DMN (13, 16, 22, 28-31). Furthermore, the RSC is one of the largest cortical regions in rodents, and there is growing evidence for functional heterogeneity along its anterior/posterior (A/P) axis (20, 29) . Based on the divergent reports in the literature, and to more accurately identify and model SN-DMN functional interactions, we directly probed the role of AI, Cg, PrL and multiple RSC subdivisions and their dynamic interactions during stimulation of the AI node of the SN.

**2. BSDS model-based estimation illustrates the power of latent space models for capturing dynamic circuit mechanisms of optogenetic stimulation**

Our analyses demonstrate that BSDS model-based estimation captures dynamic circuit mechanisms of optogenetic stimulation that are missed by conventional GLM analyses. First, conventional GLM analysis where stimulation ON and OFF blocks were directly contrasted uncovered significant response clusters in the AI, striatum, and MTL, but not in other SN and DMN nodes (**Figure S1A**). Second, gPPI analysis did not uncover significant changes in AI connectivity at the whole-brain level. Third, inspection of time-series from SN and DMN nodes revealed that the shortcomings of GLM-based approaches may be due to the complex and delayed temporal profile of fMRI signal changes elicited by optogenetic stimulation (**Figure S2**). Fourth, analysis of stimulation OFF sub-blocks following the stimulation ON block revealed delayed transition patterns (**Figure S3**) which illustrate why conventional approaches may not accurately estimate stimulation-dependent spatiotemporal changes in brain activity and connectivity. Importantly, our results highlight the importance of characterizing intermediate transition states connecting states related to stimulation ON and OFF blocks. Taken together, these results highlight the ability of latent space dynamical switching models like BSDS to identify hidden latent brain states and their spatiotemporal dynamics underlying the effects of stimulation that are missed by conventional approaches.

**3. Correspondence with the study of Mandino and colleagues (16)**

The spatial pattern of fMRI signal changes to optogenetic AI stimulation observed in our study is corroborated by recently reported findings from optogenetic AI stimulation in mice by Mandino and colleagues (16). Although differences between species (rats vs mice) and measurement modality (cerebral blood volume vs blood-oxygen-level-dependent signal) limit direct comparisons, the response areas from our GLM analysis approximate the most significant response areas reported by Mandino et al. (16), including: the AI and cortex dorsomedial to AI, a band extending from M1 through the ventral striatum, and an anterior-ventrolateral portion of MTL (**Figure S1A**). In addition, the signal profiles from individual ROIs in our study allude to responses in other locations similar to those shown by Mandino et. al. (16), such as PrL and RSC at -5.90 mm AP (**Figure S2**). Notably, CBV has been shown to be spatially (32) and functionally (33) more specific to neuronal activity than BOLD. Furthermore, Mandino et al. (16) used a considerably stronger stimulation than in the present study, including a larger bolus of optogenetic viral vector and greater stimulation light power and pulse duration despite the smaller relative volume of the mouse brain. Therefore, differences in the overall response area and regional response amplitude outside of AI could be partially explained by the relative intensity of AI stimulation between studies. In addition, as diversity in AI structural and functional connectivity has been reported along the anterior-posterior (34) and dorsal-ventral axis (13), the larger response area observed by Mandino et. al. (16) compared to the present study can also be attributed, in part, to a larger area recruited by stimulation. Nonetheless, the overall agreement between these findings provides validation for our optogenetic manipulation of AI, and points to considerable overlap in AI functional connectivity between rats and mice.

**IV. Supplementary Figures**

**Figure S1. Activation maps induced by optogenetic AI stimulation in Chronos rats.** AI stimulation increased signals in the AI, striatum, and medial temporal lobe (one sample *t*-test;
*p* < 0.005 voxel-wise threshold; *p* < 0.01, 38 voxel cluster size, family-wise cluster-correction threshold).

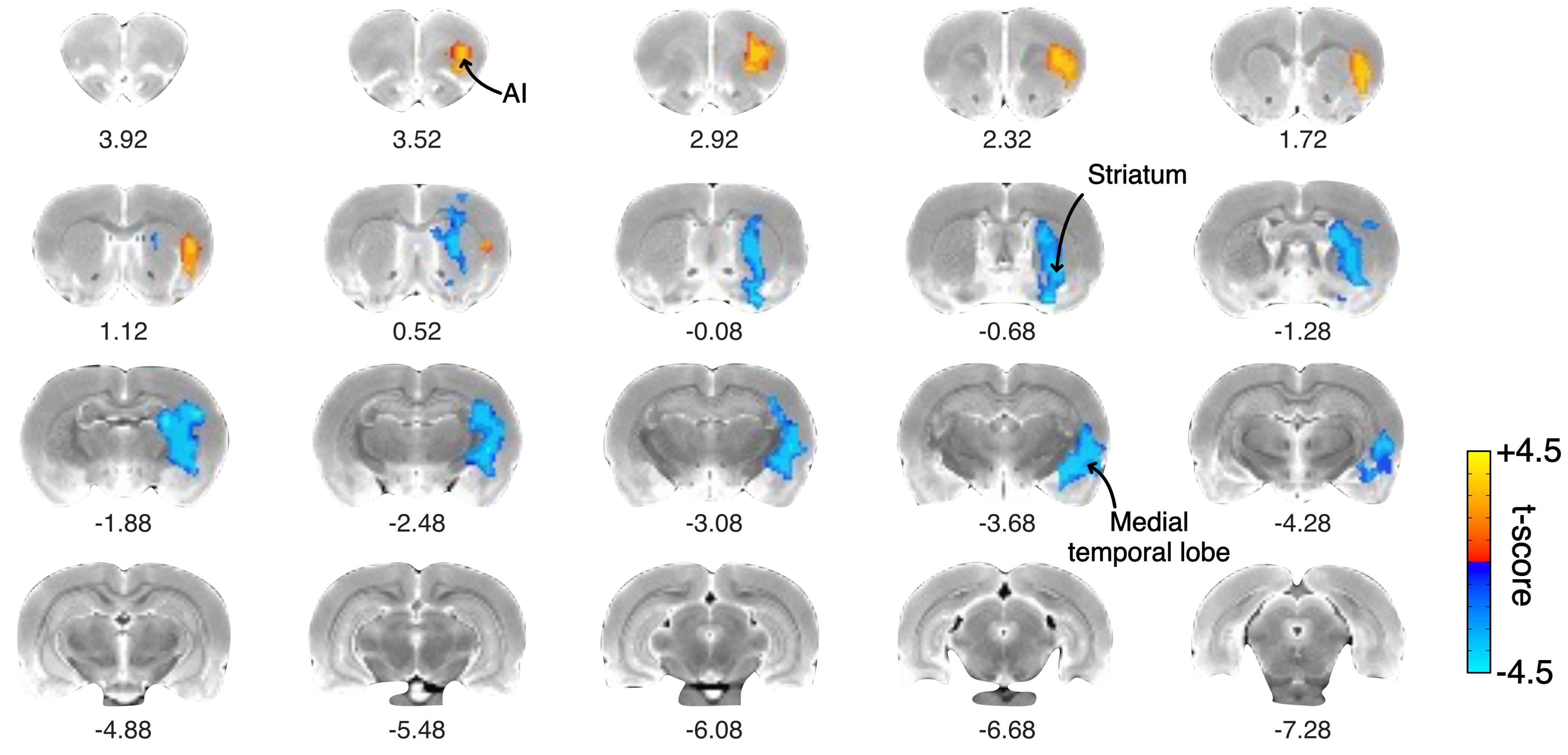

**Figure S2. Evoked responses of downstream regions to optogenetic stimulation of AI.** Averaged response of PrL, Cg, and subdivisions of RSC to optogenetic AI stimulation in Chronos rats (*N*=9). Data are presented as mean and standard error of the mean.

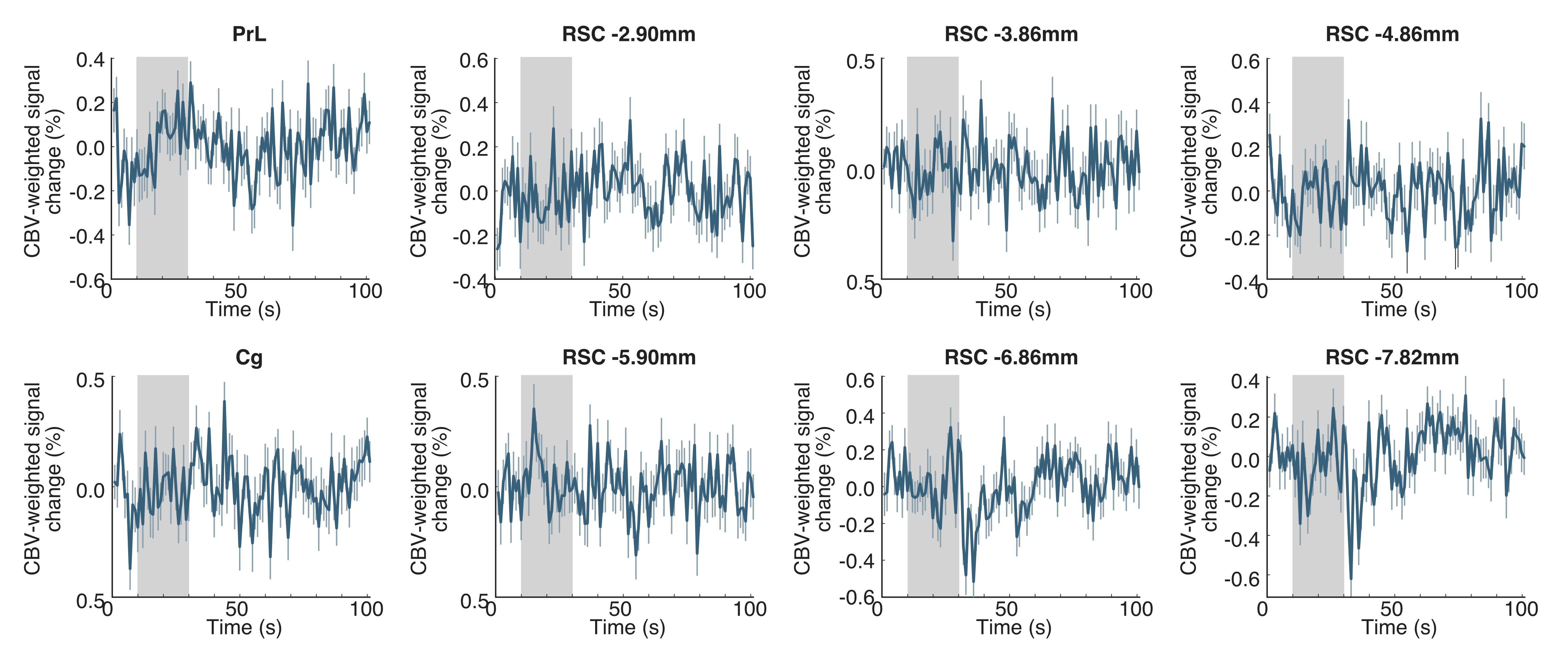

**Figure S3. Occupancy rates of latent brain states during stimulation ON and OFF blocks. (A, B)** Optogenetic stimulation protocol. We divided stimulation ON and OFF blocks into 10 s sub-blocks, respectively. **(C)** Averaged occupancy rates of latent brain states during each stimulation ON and OFF sub-block in Chronos rats (* *p* < 0.05. two-tailed *t*-test, FDR-corrected for multiple comparisons). **(D)** Averaged occupancy rates of latent brain states during each stimulation ON and OFF sub-block in EYFP controls (* *p* < 0.05. two-tailed *t*-test, FDR-corrected for multiple comparisons).

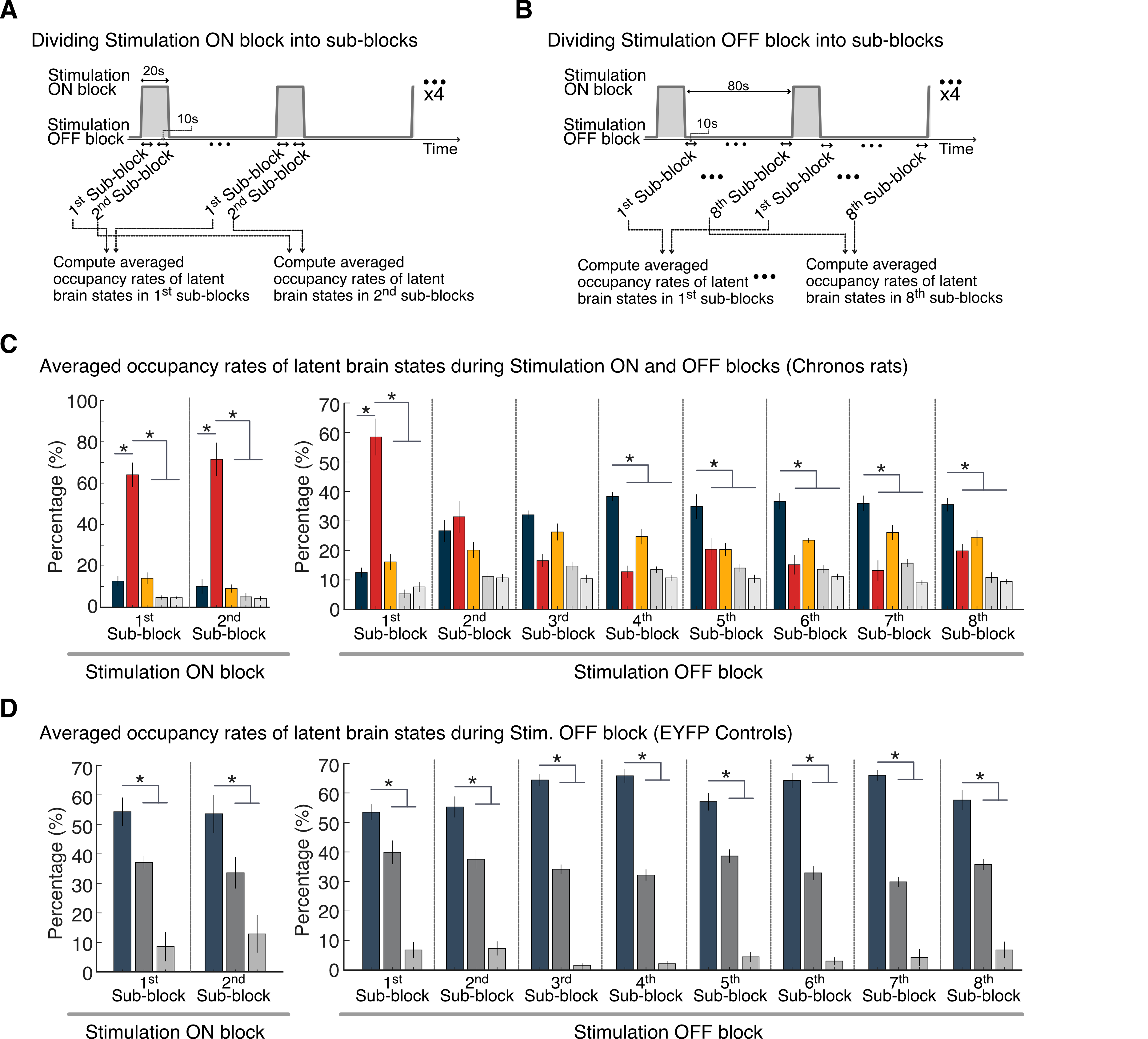

**Figure S4. Dynamic causal relationships between temporal dynamics of latent brain states (A, B)** Significant causal influences between latent brain states during the stimulation OFF and ON blocks (*p* < 0.05, two-tailed *t*-test, FDR-corrected). Red cells indicate positive influences (i.e., activation) and blue cells indicate negative influences (i.e., inhibition). **(C)** Correlation between the AI fMRI response and the temporal profile of posterior probability of State 2 (i.e., ON state). **(D)** Illustration showing dynamic causal interactions between latent brain states during the stimulation OFF and ON blocks.

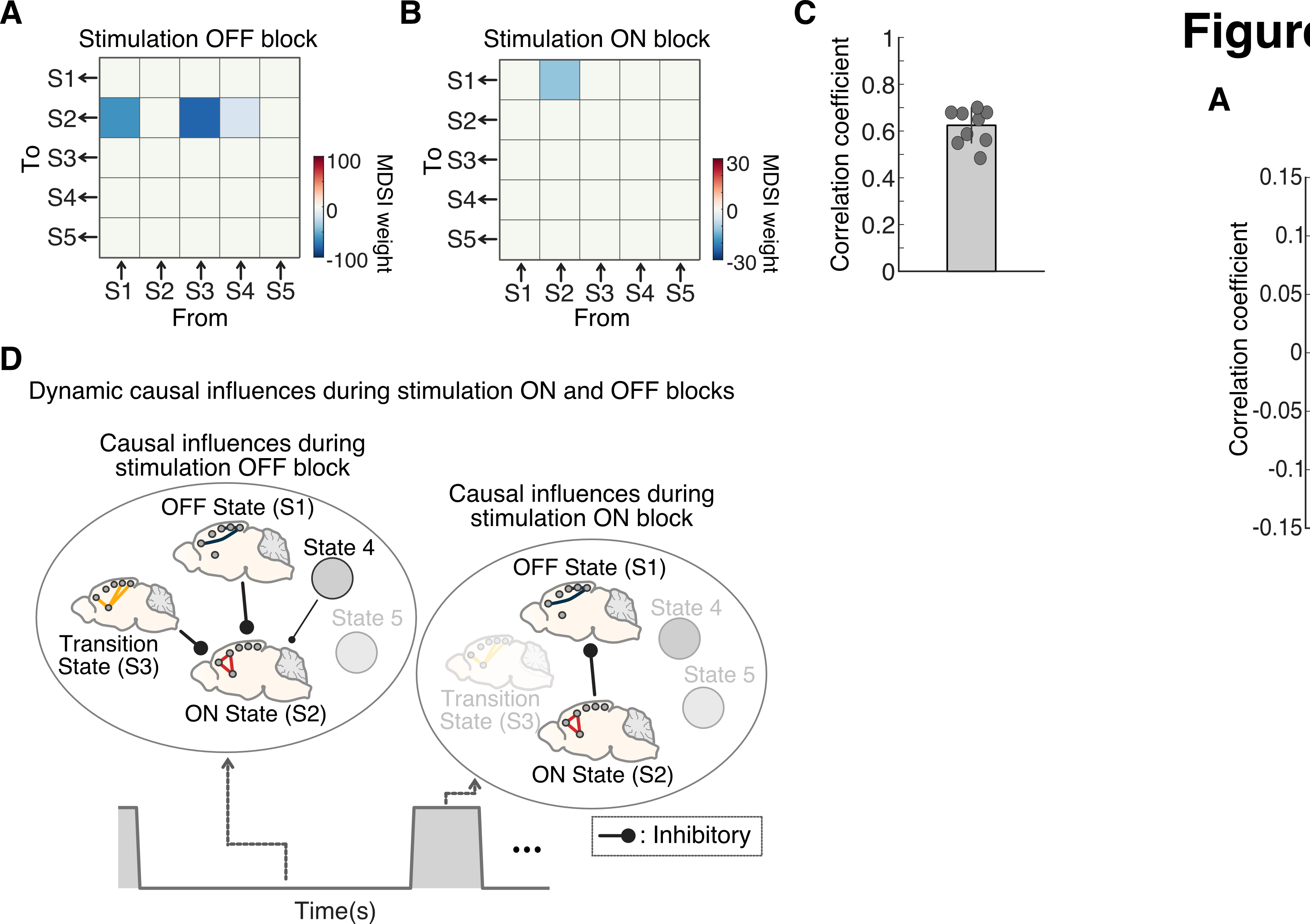

**Figure S5. Key steps in functional connectivity analysis of time-series data corresponding to BSDS-derived ON and OFF states**.

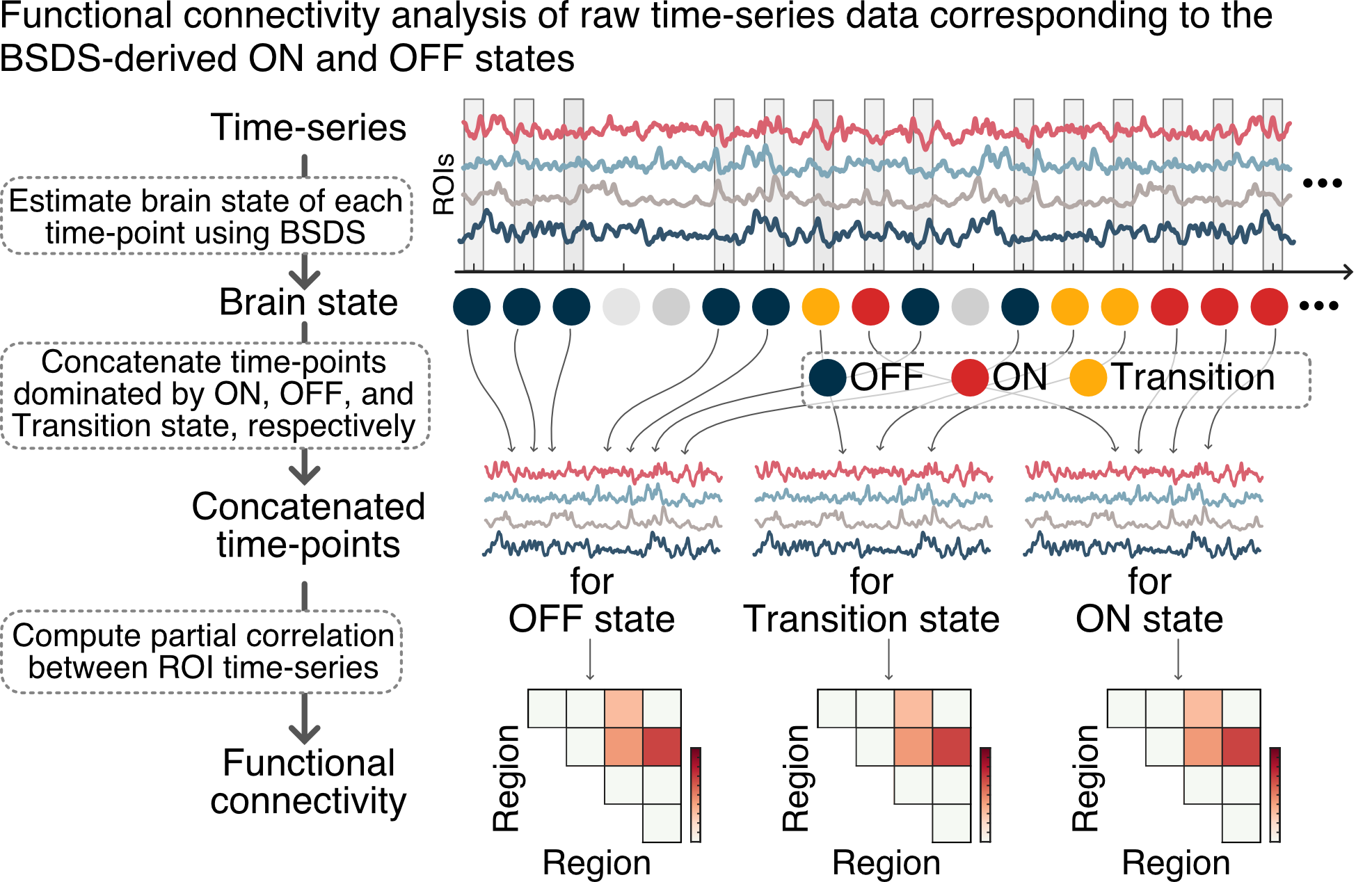

**Figure S6. Inter-regional functional connectivity using time-series data from BSDS-derived ON and OFF states. (A-C)** Functional connectivity changes between AI, PrL, and posterior RSC (-6.86mm AP) and all the other ROIs induced by optogenetic stimulation of the AI in Chronos rats. These results converge on and validate functional connectivity changes reported in Figures 4D-F. **p* < 0.05, two-tailed *t*-test, FDR-corrected.

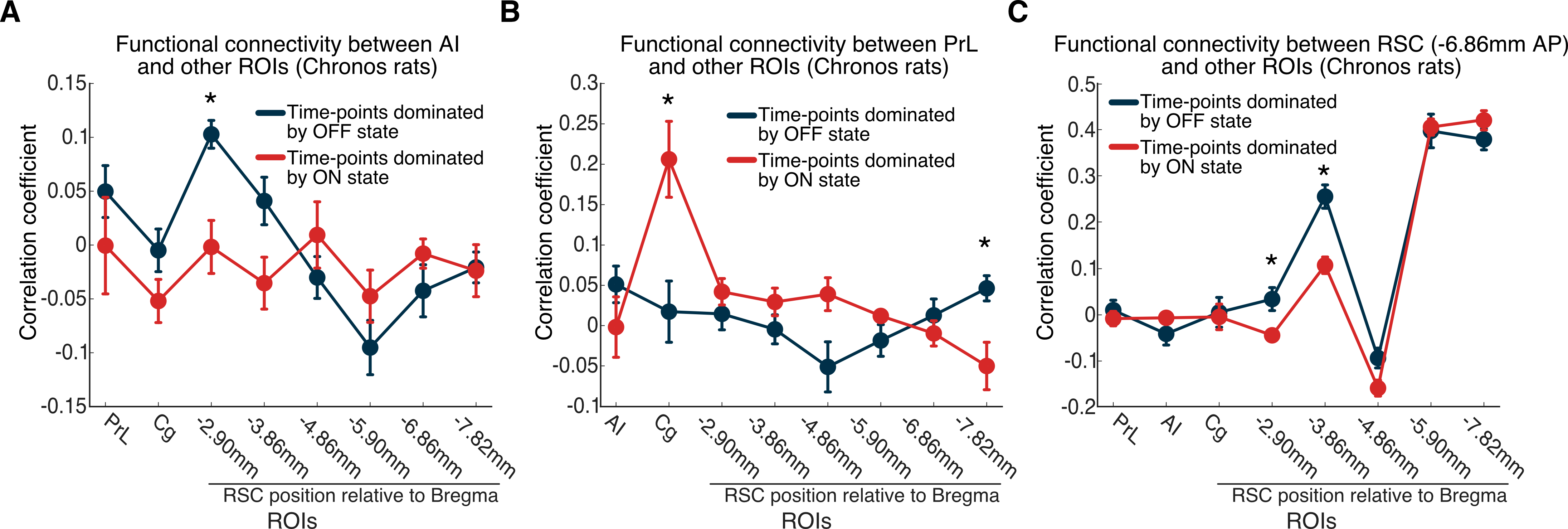

**Figure S7. State switching properties of State 3 during OFF stimulation blocks and stimulation boundaries. (A)** Schematic illustrating examination of state switching matrices during different periods of the experiment. **(B)** Occupancy rates of latent brain states during OFF stimulation blocks and stimulation boundaries. **(C-E)** State switching matrices identified state transitions during OFF stimulation blocks, OFF🡪ON stimulation boundaries and ON🡪OFF stimulation boundaries, respectively. **(F-H)** Likely transition path during OFF stimulation blocks, OFF🡪ON stimulation boundaries and ON🡪OFF stimulation boundaries, respectively, based on the state switching probabilities between states in panel **C-E**.

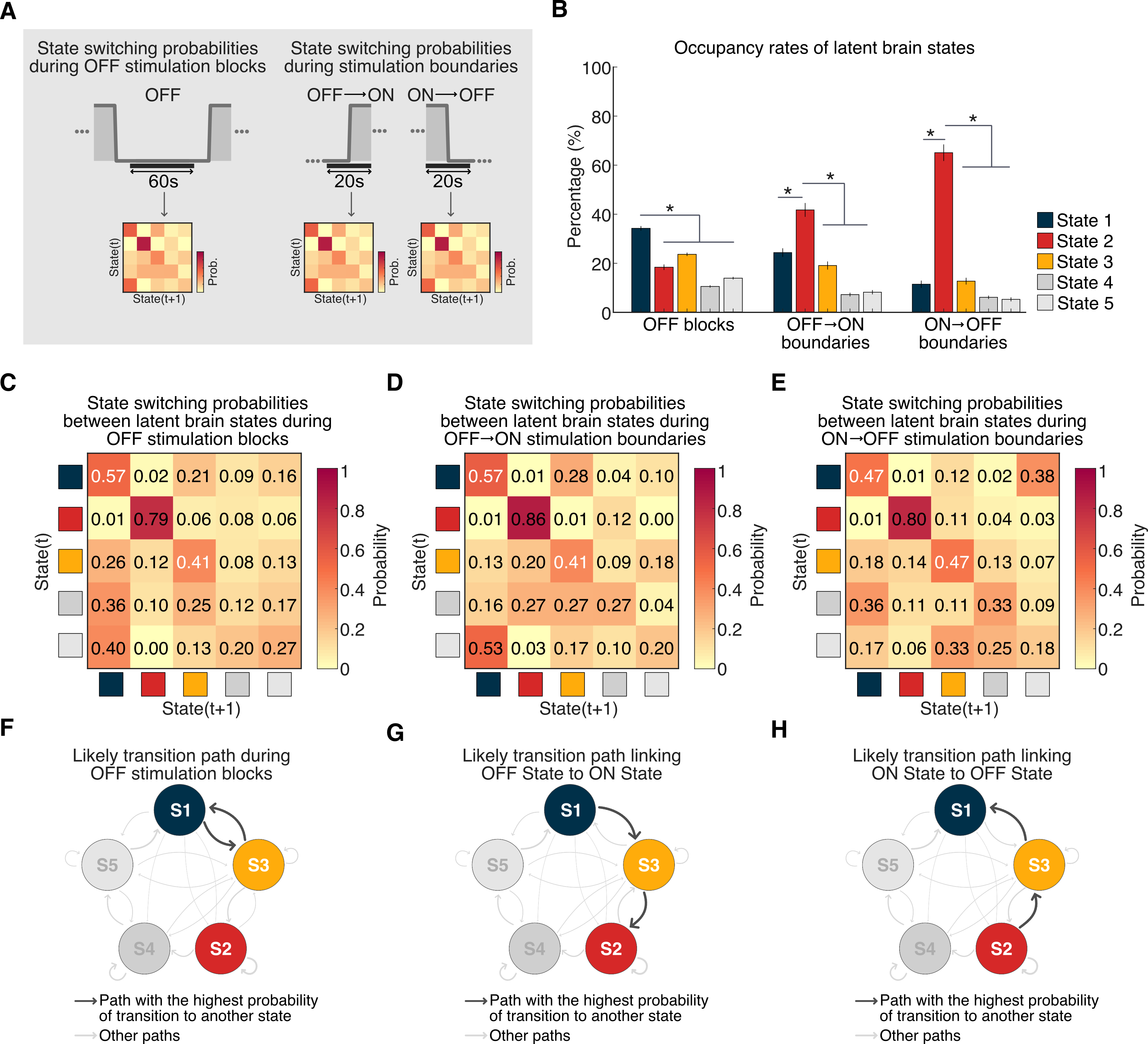

**Figure S8. Replication of dynamic functional connectivity changes during the Transition compared to the ON and OFF states. (A)** Comparison of functional connectivity between Transition and OFF states. **(B)** Comparison of functional connectivity between ON and Transition states. These results converge on and validate functional connectivity changes reported in **Figures 6C and 6F.**

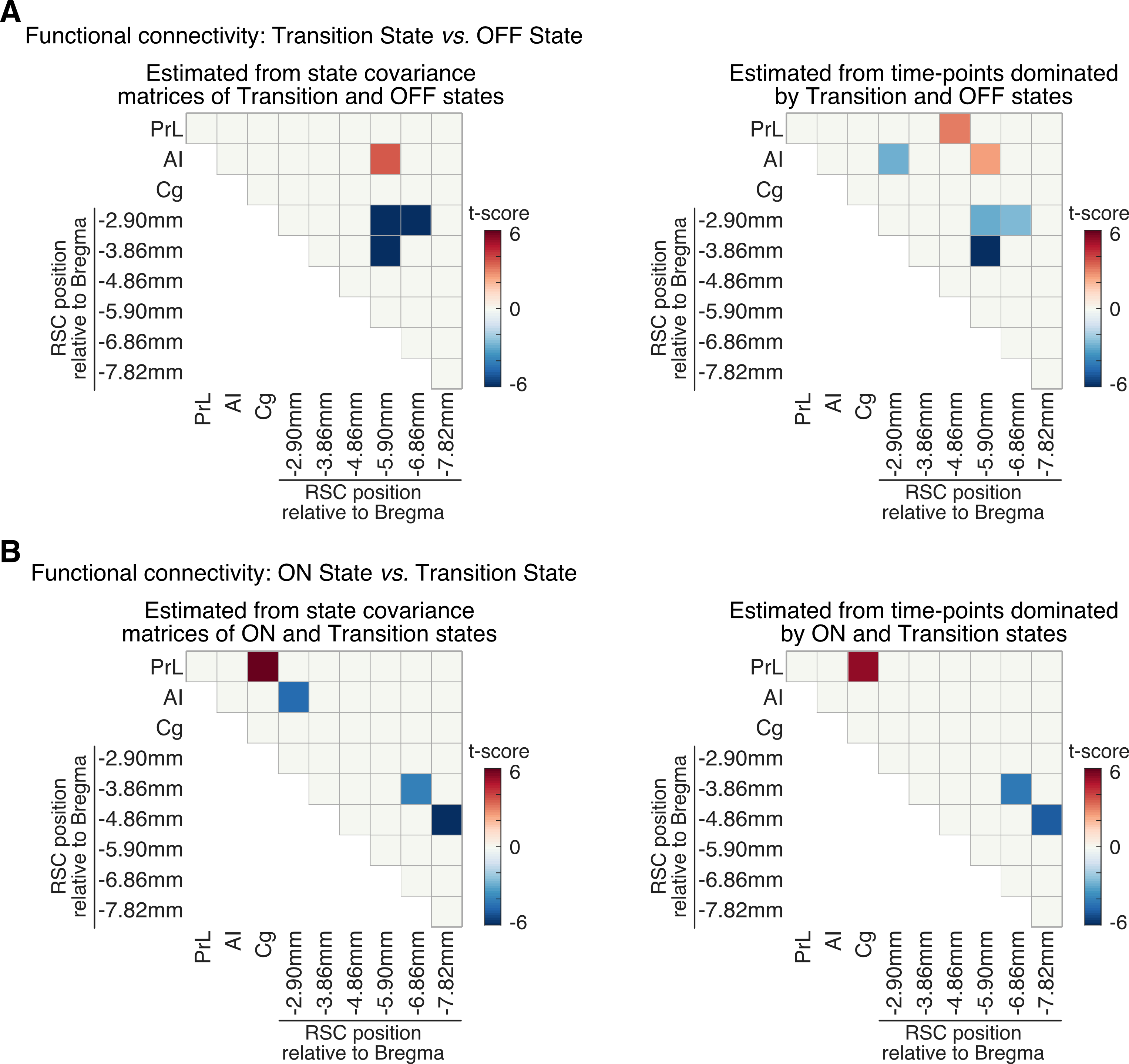

**Figure S9.** **Replication of findings using extended SN and DMN ROIs. (A)** Regions of interests used in the extended model. **(B)** Correlation of posterior probabilities of latent brain states estimated from the original 9 ROI model and 11 ROI extended model. **(C)** Averaged time-varying posterior probabilities of the brain states identified in Chronos rats by the BSDS model across the AI stimulation protocol (top). Temporal evolution of the brain states identified in Chronos rats (bottom). **(D)** Occupancy rates of the latent brain states in Chronos rats. Data are represented as mean and standard error of the mean (* *p* < 0.05, two-tailed *t*-test, FDR-corrected). **(E)** Specific links that showed significant differences in functional connectivity between the stimulation ON and OFF states (all *ps* < 0.05, two-tailed *t*-test, FDR-corrected).

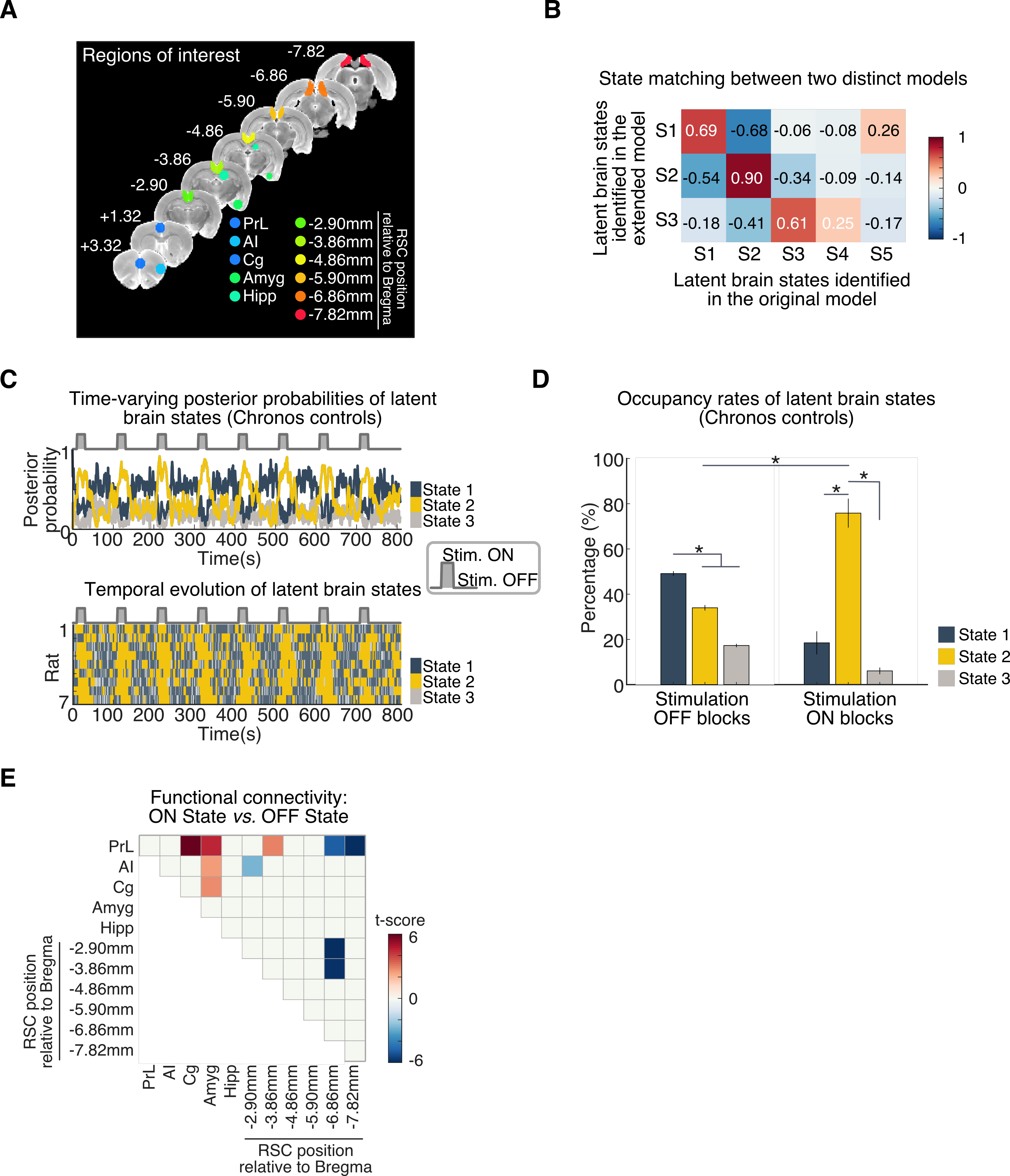

**Figure S10.** **Replication of findings using SN and DMN nodes and additional striatum-MTL ROIs. (A)** Regions of interests used in the extended model. **(B)** Correlation of posterior probabilities of latent brain states estimated from the original 9 ROI model and 11 ROI extended model. **(C)** Averaged time-varying posterior probabilities of the brain states identified in Chronos rats by the BSDS model across the AI stimulation protocol (top). Temporal evolution of the brain states identified in Chronos rats (bottom). **(D)** Occupancy rates of the latent brain states in Chronos rats. Data are represented as mean and standard error of the mean (* *p* < 0.05, two-tailed *t*-test, FDR-corrected). **(E)** Specific links that showed significant differences in functional connectivity between the stimulation ON and OFF states (all *ps* < 0.05, two-tailed *t*-test, FDR-corrected).

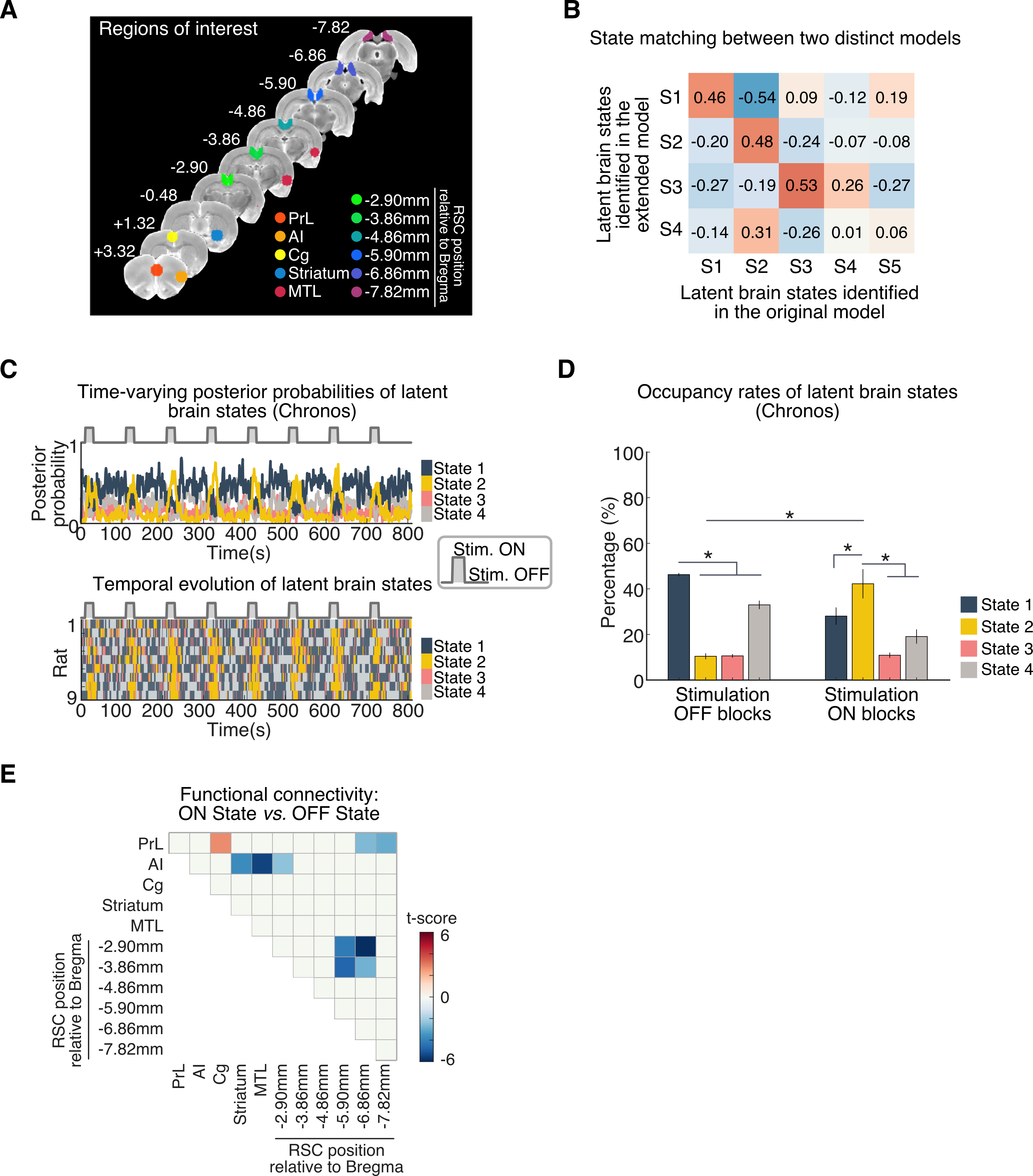

**Figure S11.** **Control analysis using ROIs located outside the DMN and SN. (A)** Regions of interests (ROIs) used in control analysis. **(B)** Averaged time-varying posterior probabilities of the brain states identified in Chronos rats by the BSDS model across the AI stimulation protocol (top). Temporal evolution of the brain states identified in Chronos rats (bottom). **(C)** Occupancy rates of the latent brain states in Chronos rats. State 1 dominated both stimulation ON and stimulation OFF blocks. Data are represented as mean and standard error of the mean.

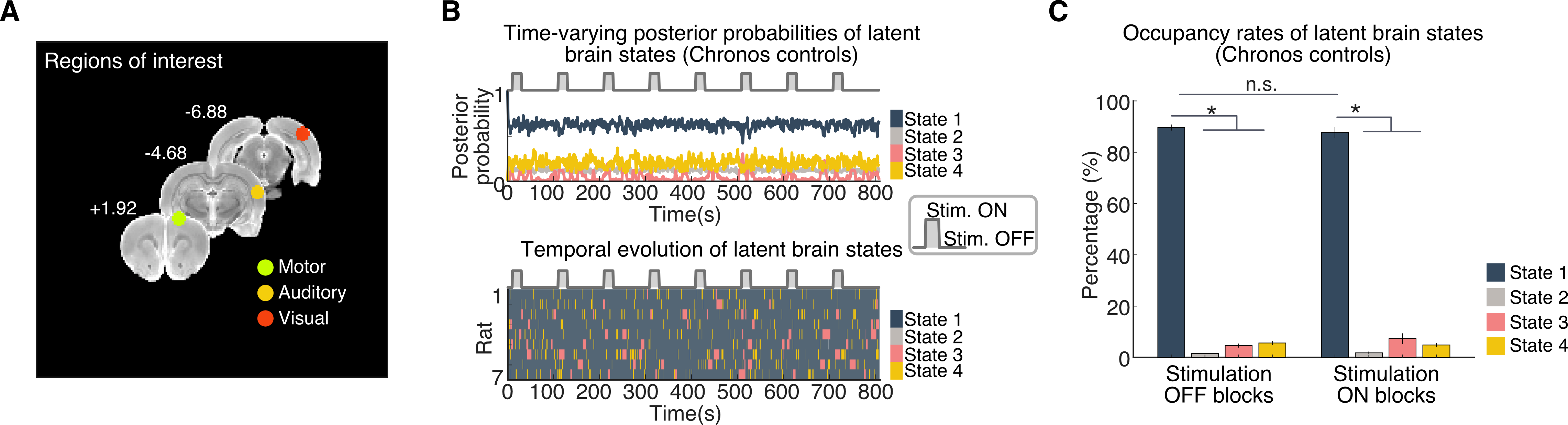
